# The cyclic nitroxide antioxidant 4-methoxy-TEMPO decreases mycobacterial burden *in vivo* through host and bacterial targets

**DOI:** 10.1101/464586

**Authors:** Harrison D Black, Wenbo Xu, Elinor Hortle, Sonia I Roberston, Warwick J Britton, Amandeep Kaur, Elizabeth J New, Paul K Witting, Belal Chami, Stefan H Oehlers

## Abstract

Tuberculosis is a chronic inflammatory disease caused by persistent infection with *Mycobacterium tuberculosis*. The rise of antibiotic resistant strains necessitates the design of novel treatments. Recent evidence shows that not only is *M. tuberculosis* highly resistant to oxidative killing, it also co-opts host oxidant production to induce phagocyte death facilitating bacterial dissemination. We have targeted this redox environment with the cyclic nitroxide derivative 4-methoxy-TEMPO (MetT) in the zebrafish-*M. marinum* infection model. MetT inhibited the production of mitochondrial ROS and decreased infection-induced cell death to aid containment of infection. We identify a second mechanism of action whereby stress conditions, including hypoxia, found in the infection microenvironment appear to sensitise *M. marinum* to killing by MetT both *in vitro* and *in vivo*. Together, our study demonstrates MetT inhibited the growth and dissemination of *M. marinum* through host and bacterial targets.

## 1. Introduction

Tuberculosis (TB) is a communicable disease of global health concern. TB is caused by persistent infection with *Mycobacteria tuberculosis (Mtb)*, an intracellular bacterial pathogen that can evade the immune response and persist in even immunocompetent patients for life. The prevalence of antibiotic resistance in circulating *Mtb* strains is rising, highlighting the need for novel anti-infective and host tissue-sparing strategies [1].

The pathogenesis of *Mtb* infection is a function of the complex interaction between the virulence of *Mtb* and the host immune response [2]. Within macrophages, *Mtb* faces many mechanisms of killing including lysosomal acidification of the phagosome and oxidative killing [3]. Cytokine signalling from IL-1β and TNFα, and pattern recognition signalling induce the expression of phagocytic NADPH oxidase (NOX) and inducible nitric oxide synthase (iNOS) in mycobacteria-infected macrophages [3, 4]. NOX mediates the production of superoxide ions (O_2_^-^), react further with water to form highly reactive species such as hydrogen peroxide (H_2_O_2_) and hydroxyl radicals (•OH), which are key examples of reactive oxygen species (ROS). ROS are highly effective microbicidal molecules, but pathogenic mycobacteria are highly resistant to oxidative mediated killing through expression of catalase, superoxide dismutase, and peroxiredoxin enzymes that detoxify ROS to benign products [3].

Inflammation stimulates expression of NOX and NOS, and the production of ROS in the mitochondria [5]. Disruption of the tightly regulated balance between oxidants and antioxidants results in pathological oxidation of proteins, lipids, and nucleic acid causing oxidative stress and cell death [6]. Evidence suggests that pathogenic mycobacteria co-opt the host immune response to facilitate a growth-permissive environment, through the modulation of host oxidant production. Roca and Ramakrishnan showed that mitochondrial superoxide was upregulated during mycobacterial infection caused by signalling from TNF through the RIP1/RIP3 pathway favouring necroptotic cell death and the detrimental release of intracellular mycobacteria [7]. Further, they demonstrated oxidant scavenging molecules, including the cyclic nitroxide TEMPOL, were able to inhibit the growth of mycobacteria and cell death in conditions of TNF overexpression but not in wild type animals [7].

Dallenga *et al*. also demonstrated that host oxidant production benefited pathogen replication using an *in vitro* macrophage and neutrophil co-culturing model to show that mycobacterial ESAT-6 induced neutrophil ROS production and neutrophil necrosis [8]. Necrotic neutrophil debris were then phagocytosed by macrophages, which also underwent necrotic cell death thus promoting the growth of *Mtb*. Inhibition of ROS production by neutrophil myeloperoxidase inhibited necrosis and bacterial growth suggesting that oxidant production by this class of leukocytes may be a target for therapeutic development.

There is also evidence that *Mtb* seeks to inhibit host antioxidant capacity. Human patients show increased levels of systemic oxidative damage, and a reduced systemic antioxidant capacity [9]. Palanisamy *et al*. recapitulated these findings in a guinea pig *Mtb* infection model, and showed that treatment with the oxidant scavenger N-acetyl cysteine (NAC), partially ablated the reduced antioxidant capacity and decreased markers of oxidative damage at the site of the lesion [10]. They also showed that NAC-treated animals had fewer necrotic granulomas.

Cyclic nitroxides are low molecular weight stable radicals that have potent antioxidant activity *in vitro* and *in vivo*, and have been cleared for human clinical use [11]. They readily cross cell membranes, and diffuse into tissues [12, 13]. They afford extensive protection to cells when toxicity is mediated by the production of ROS and RNS [14]. 4-Methoxy-2,2,6,6-tetramethylpiperidine 1-oxyl, shortened to MetT, is a stable cyclic nitroxide derivative based on TEMPO, with a substitution at position 4 for a methoxy group [15]. The complete characterisation of the functional difference of this substitution to other nitroxides is lacking in the literature, but MetT appears to have similar activity to other cyclic nitroxides [16].

Given the serious issues with the current pathogen directed treatment paradigm in TB, and the evidence that host oxidant production is implicated in the progression of *Mtb* infection, there is a basis for investigating cyclic nitroxide antioxidants as host directed therapies for TB. Here we investigate the utility and mechanisms of action of MetT as a model powerful antioxidant therapy for TB using the zebrafish-*M. marinum* infection model.

## 2. Materials and Methods

### 2. 1. Zebrafish husbandry methods

Adult zebrafish (*Danio rerio*) were maintained at the Garvan Institute Biological Testing Facility and eggs were collected by natural spawning (St Vincent’s Hospital AEC Approval 1511). Lines used in this study: TAB strain wild types, *Tg*(*lyzC:DsRed*^*nz*50^), *Tg*(*mfap4:TdTomato*^*xt*12^), *Tg*(*mpeg1:tomato-caax*^*xt*3^), Durif mutant in mpx gene (*drf*^*g*l8^) *Tg*(*mpx:EGFP*^*i*114^). All experiments and procedures were completed in accordance with Sydney Local Health District Animal Welfare Committee for zebrafish embryo research.

### 2. 2. Microinjection of Mycobacterium marinum

Infection was carried out as previously described [17].

### 2. 3. Drug treatments

Drugs were added directly to zebrafish media (E3 supplemented with PTU to inhibit pigmentation) immediately after infection unless otherwise indicated. Media containing drugs was replenished every second day for the duration of the experiment. Zebrafish experiments used final concentrations of 1 mM MetT, 3 µM diphenyleneiodonium (DPI) and 50 µg/ml dexamethasone.

### 2. 4. Imaging embryos

Epifluorescence microscopy was performed using a Leica M205FA fluorescence stereomicroscope or Leica DM600B light and fluorescence microscope. Confocal microscopy was performed with a Leica SP5 inverted confocal microscope. Images were processed initially with Leica native LAS software then with ImageJ (NIH) and Photoshop CS6 (Adobe).

### 2. 5. Dihydrorhodamine staining

Biological oxidants were identified *in vivo* with live embryos treated with 500 μM dihydrorhodamine-123 (DHR; Sigma) and incubated for 30 minutes in dark conditions. Embryos were extensively washed prior to imaging.

### 2. 6. FRR2 mitochondrial oxidant probe staining

Mitochondrial ROS during mycobacterial infection was quantified using the flavin-rhodamine redox sensor 2 (FRR2) probe with confocal microscopy [18]. Embryos were treated with 100 μM FRR2 for 2 hours in dark conditions at 28°C, extensively washed and mounted in 1% (w/v) low melting point agarose for confocal imaging.

Katushka fluorescent protein (expressed by *M. marinum*) was excited in one scan, with a 561 nm laser set to and detector set to 700-800 nm. The mitochondrial probe FFR2 was excited with another scan with the 488 nm argon laser and the detector for red emission set to 560-630 nm.

### 2. 7. TUNEL assay

Cell death was quantified with a Click-iT™ Plus TUNEL Assay for In Situ Apoptosis Detection, Alexa Fluor™ 488 dye kit (ThermoFisher) according to manufacturer’s directions. Briefly, embryos were fixed in 10% v/v neutral buffered formalin (NBF) overnight at 4 ̊C, washed in PBS then permeabilised by digestion with 10 µg/mL proteinase K for 30 min before post fixing in 10% NBF at 37 °C for 5 min. Embryos were preincubated in Tdt reaction buffer for 10 min at 37 °C then for a further 30 min at 37 °C with Tdt reaction buffer containing Tdt enzyme and EdUTP. Following Tdt reaction embryos were rinsed in PBS and incorporated EdUTP was detected with the Click-iT™ plus TUNEL reaction cocktail for 30 min at 37 °C in the dark, which incorporates Alexa Fluor 488 fluorescent marker to the EdUTP by Click chemistry. Embryos were extensively rinsed in PBS prior to imaging.

### 2. 8. Image processing and analysis

Infection burden and fluorescent probe staining were measured as the number of pixels above background per embryo in ImageJ using binary thresholding of single florescence channel images and the Analyze Particles function. Fluorescent probe intensity was measured as the maximum signal using the ImageJ Measure function.

### 2.9. Quantitative PCR

Pooled embryos (10-40) were homogenised with a 21-gauge syringe in Trizol reagent (Thermo Fisher Scientific) for total RNA extraction. cDNA was reverse transcribed using the High-Capacity cDNA Reverse Transcription Kit (Thermo Fisher Scientific). qPCR was carried out using PowerUP SYBR Green Mastermix (Thermo Fisher Scientific) on an Aglient

Technologies Mx300P thermal cycler. PCR cycling conditions were 95°C for 10 minutes, then 40 cycles of 95°C for 15 seconds and 60 degrees for 1 minute, followed by a melt curve between 55°C and 95°C with 1.5°C/second ramp rate. Primer pairs 5’-3’ were: *il1b* ATCAAACCCCAATCCACAGAGT and GGCACTGAAGACACCACGTT, il8 TGTTTTCCTGGCATTTCTGACC and TTTACAGTGTGGGCTTGGAGGG, ef1a TGCCTTCGTCCCAATTTCAG and TACCCTCCTTGCGCTCAATC, tnf GTTTATCAGACAACCGTGGCCA and GATGTTCTCTGTTGGGTTTCTGAC.

### 2.10. Mycobacterial *in vitro* culture and CFU measurements

*M. marinum*-TdTomato or *M. marinum* Δesx1 *pCherry/peredox* was grown in 7H9 Mycobacterial media (Difco Middlebrook), enriched with 10% w/v OADC and 50 µg/mL hygromycin, and cultured at 28°C without agitation with drugs at final concentrations indicated. Samples of the broth culture were serially diluted and plated on 7H10 Mycobacterial agar (Difco). Plates were incubated for at least 7 days at 28°C and CFU were manually counted.

### 2.11. Hypoxic culture

Bacterial cultures or zebrafish embryos were co-incubated with an Oxoid™ CampyGen™ 2.5 L Sachet (Thermo Fisher Scientific) and maintained in either a 2.5 L (bacterial cultures) or 5 L (zebrafish) airtight container for an expected oxygen concentration of 5% or 10% respectively.

### 2.12. Statistical analysis

Data are presented as means ± standard deviation. Statistical tests as indicated were carried out in Prism 7 (Graph Pad) using T-tests for experiments with only two groups and ANOVA for multiple comparisons using Tukey post hoc test. Significance is indicated as: P < 0.05 *, P < 0.01 **, P < 0.001 ***.

## 3. Results

### 3. 1. Macrophages and neutrophils are both prolific producers of oxidants during mycobacterial infection of zebrafish

To investigate the sources of host oxidant production, macrophage reporter embryos *Tg*(*mfap4:tdTomato*)^*xt*12^ and neutrophil reporter embryos *Tg*(*lyzC:DsRed*)^*nz*50^ were infected with fluorescent *M. marinum* and stained with the probe dihydrorhodamine 123 (DHR).

Imaging of rhoadamine123^+^ fluorescence identified colocalisation of *M. marinum*-Katushka and macrophages in *Tg*(*mfap4:tdTomato*)^*xt*12^ embryos (Fig 1A). Many of the infected macrophages also showed colocalisation of staining with rhodamine123, indicating extensive production of oxidants within those macrophages, as anticipated from previous work by Roca and Ramakrishnan [19].

**Figure 1:**
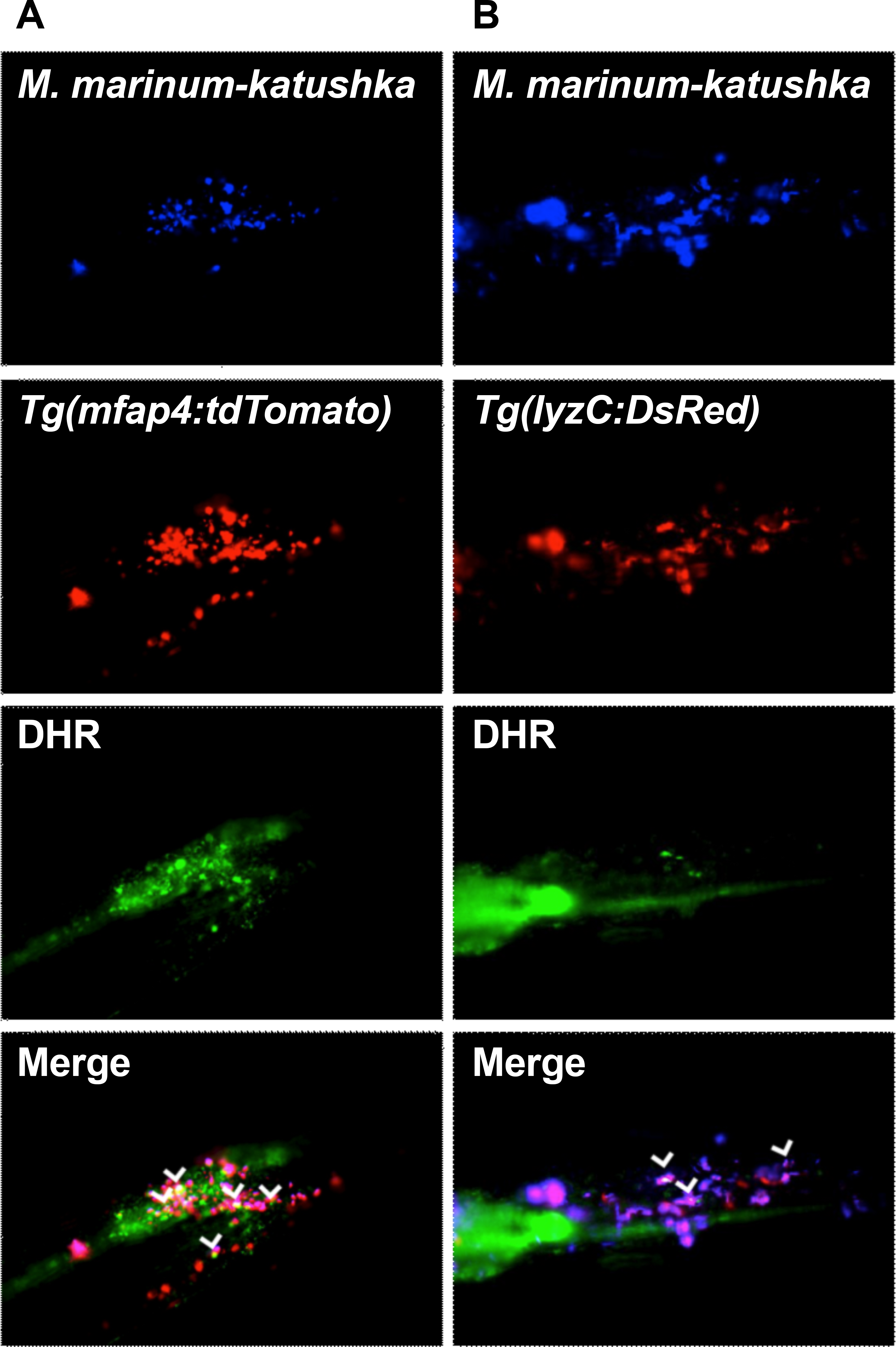
Macrophages and neutrophils produce ROS during mycobacterial infection in zebrafish. (A) Representative images of DHR-stained ROS production by *mfap4*-expressing macrophages in a zebrafish infected with *M. marinum*-katushka. (B) Representative images of DHR-stained ROS production by *lyzC*-expressing neutrophils in a zebrafish infected with *M. marinum*-katushka. Arrowheads indicate fluorescent leukocytes with internalised *M. marinum* producing ROS.

Similarly, fluorescent imaging of *M. marinum*-infected *Tg*(*lyzC:dsred*)^*nz*50^ neutrophil reporter embryos revealed colocalisation of *M. marinum*-Katushka and neutrophils, however to a much lesser extent than *M. marinum* and macrophages (Fig 1B). DHR staining revealed extensive colocalisation of infected *lyzC:dsred* neutrophils and rhodamine indicating production of oxidants within those neutrophils at the site of infection.

### 3. 2. The cyclic nitroxide antioxidant 4-methoxyTEMPO decreases mycobacterial burden in the zebrafish embryo

Next we examined the potential for the cyclic nitroxide antioxidant 4-methoxy-TEMPO (MetT) to inhibit mycobacterial growth and dissemination *in vivo*. Zebrafish embryos were infected with fluorescent *M. marinum*-TdTomato and imaged for bacterial burden at 3 and 5 days post infection (dpi) with drug treatment initiated immediately after infection (Fig 2A and 2B).

**Figure 2:**
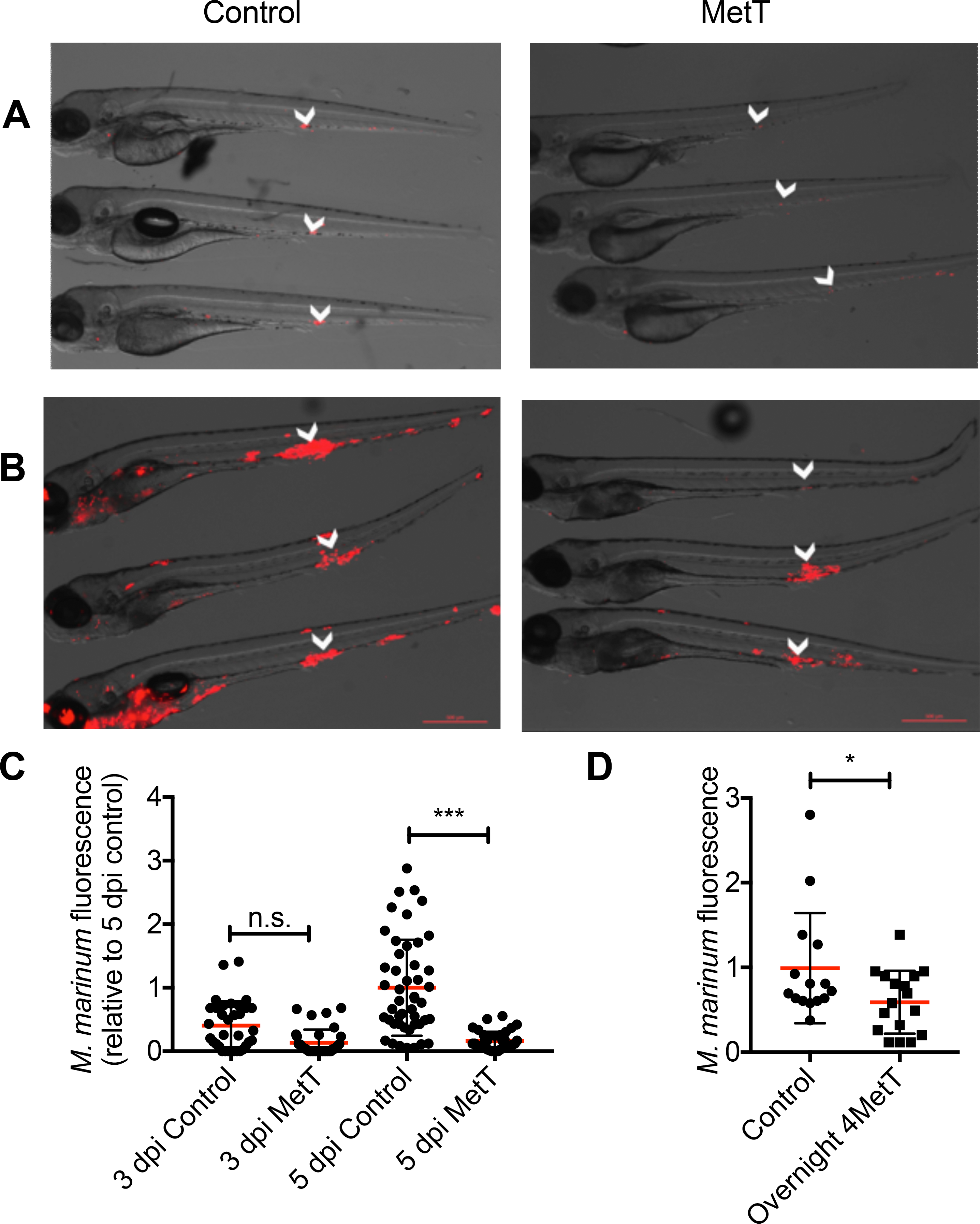
MetT treatment reduces mycobacterial burden *in vivo*. (A) Representative images of 3 dpi zebrafish embryos infected with *M. marinum*-tomato and treated with MetT. Arrowheads indicate site of primary infection. (B) Representative images of 5 dpi zebrafish embryos infected with *M. marinum*-tomato and treated with MetT. Arrowheads indicate site of primary infection. (C) Quantification of bacterial burden by fluorescent pixel count in 3 and 5 dpi zebrafish embryos treated with MetT. (D) Quantification of bacterial burden by fluorescent pixel count in 5 dpi zebrafish embryos treated overnight with MetT at 4 dpi.

In untreated control embryos, widespread dissemination of bacteria was observed from the site of infection at the caudal vein and deeper tissues throughout the head and pericardium. However, the bacteria were much less disseminated and relatively more confined to the caudal vein region in the MetT treated embryos, with only very small granulomas forming outside the site of initial infection (Fig 2B).

A fluorescent pixel count method was used to estimate bacterial burden in these infected embryos (Figure 2C). At 3 dpi, MetT-treated embryos had similar bacterial burden compared with untreated infected control embryos. However, at 5 dpi, MetT-treated embryos had significantly less bacterial burden than control embryos.

We next determined whether MetT treatment would be active against an established infection by waiting until 4 dpi to start treatment. Analysis of bacterial burden at 5 dpi in this experimental model again revealed less bacteria in the treated group than in the control group demonstrating a relatively late window of treatment efficacy (Fig 2D).

### 3.3. MetT reduces mitochondrial oxidants within the granuloma

Cyclic nitroxides are known to decrease the accumulation of oxidants in biological environments through multiple mechanisms including direct radical scavenging and through acting as a superoxide dismutatse mimetic [14, 15, 20]. Given that phagosomal oxidants are associated with host protection and bactericidal action, we hypothesised that MetT may act on a different component to inhibit bacterial growth *in vivo*. Mitochondrial ROS is upregulated during mycobacterial infection and is associated with host-cell death and the dissemination of the infection [7, 8], and we hypothesized that the cyclic nitroxide may regulate this source of ROS.

We visualised mitochondrial ROS during mycobacterial infection using confocal microscopy of zebrafish embryos stained with the flavin-rhodamine redox sensor 2 (FRR2) probe to investigate the potential of MetT to regulate mitochondrial ROS production [18]. Oxidised FRR2 exhibits a strong red fluorescence that we could quantify from confocal microscopy. Zebrafish embryos were infected with *M. marinum*-katushka and stained with FRR2 probe at 5 dpi. FRR2 fluorescence extensively localised with areas of *M. marinum*, indicating mitochondrial ROS production at the site of infection (white arrows in Fig 3A), while the adjacent non-infected tissue did not show evidence of FRR2 fluorescence (Fig 3A).

**Figure 3:**
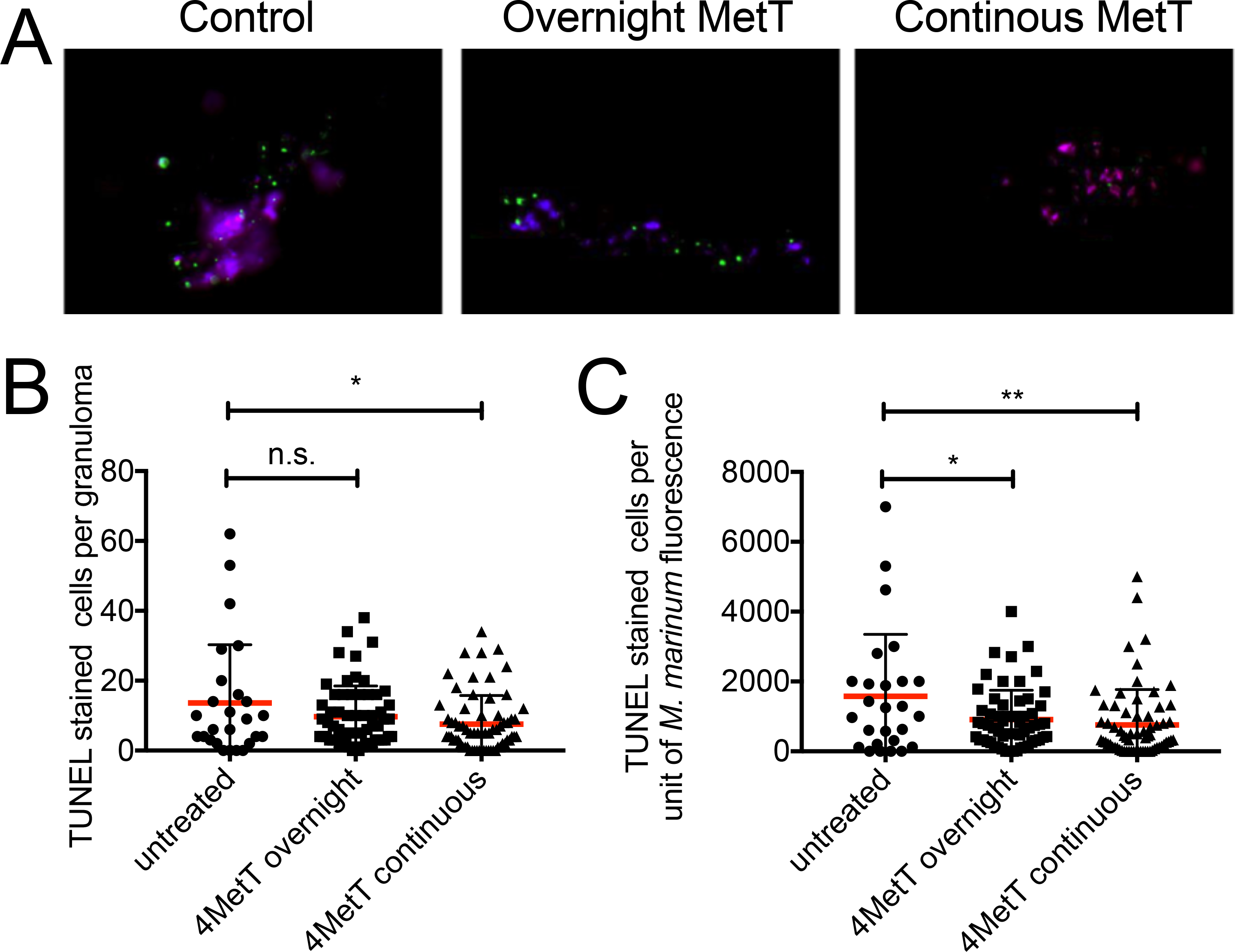
MetT treatment reduces mitochondrial ROS production in mycobacterial granulomas. (A) Representative images of 5 dpi zebrafish embryos infected with *M. marinum*-katushka (blue) stained with FRR2 (red, and purple colocalisation) for mitochondrial ROS production. (B) Representative images of 5 dpi zebrafish embryos infected with *M. marinum*-katushka (blue) stained with FRR2 (red, and purple colocalisation) for mitochondrial ROS production and treated with MetT. (C) Quantification of mitochondrial ROS production as a ratio of bacterial content per granuloma in 5 dpi zebrafish embryos treated with MetT.

Quantification of infection-induced FRR2 oxidised from fluorescence above background fluorescence revealed a lower intensity of red fluorescent FRR2 per granuloma in MetT-treated embryos (Fig 3B). Given that the lower bacterial burden of MetT-treated embryos could be expected to correlate with decreased mitochondrial ROS production, we next calculated the proportional area of FRR2 fluorescence per unit of mycobacteria in each granuloma. Again, MetT-treated embryos showed reduced area of oxidised FRR2 probe per unit of mycobacteria per granuloma compared to control embryos that were not treated with MetT (Fig 3C).

### 3.4. MetT reduces cell death in the mycobacterial granuloma

Having shown that MetT acts to reduce mitochondrial ROS in the granuloma and inhibits the growth of *M. marinum* in the zebrafish embryo we next sought to determine the effect of MetT on cell death in the mycobacterial granuloma. Terminal deoxynucleotidyl transferase dUTP nick end labelling (TUNEL) allows for the in situ identification of cells that have undergone necroptotic and apoptotic cell death.

Embryos were infected with *M. marinum*-Katushka, and treated either continuously with MetT or overnight with MetT from late in 4 dpi and were fixed at 5 dpi for TUNEL staining (Figure 4A). Embryos that were continuously treated with MetT had significantly less TUNEL stained cells per granuloma than untreated control embryos (Fig 4B). Additionally, calculating the number of TUNEL stained cells per unit of fluorescent mycobacteria per granuloma revealed a significant reduction in cell death between both MetT treatment groups and the untreated control group (Fig 4C). These data demonstrate MetT reduces immunopathology and bacterial spread through directed inhibition of cell death in the mycobacterial granuloma.

**Figure 4:**
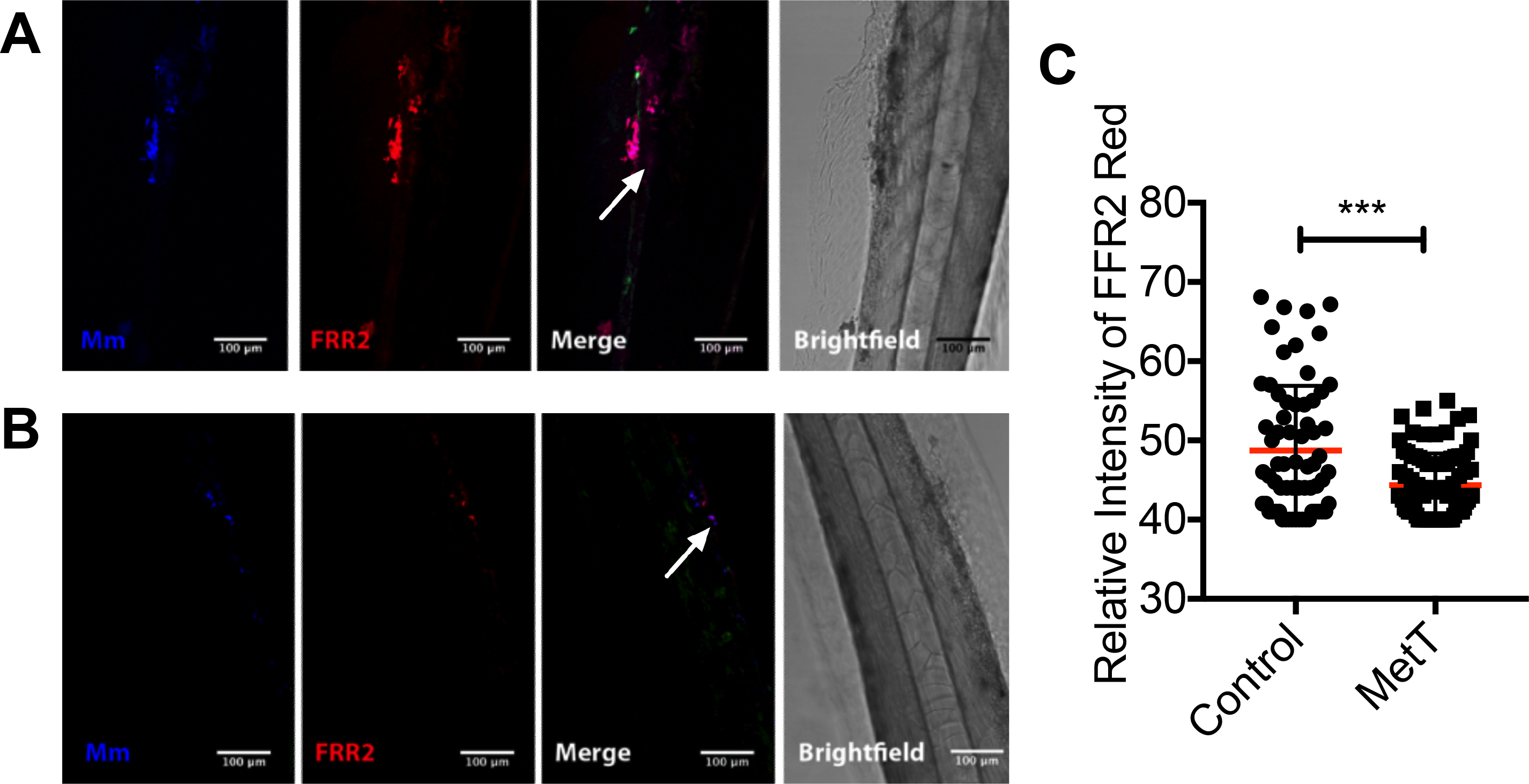
MetT treatment reduces cell death in mycobacterial granulomas. (A) Representative images of TUNEL staining for apoptotic and necroptotic cells in 5 dpi zebrafish embryos infected with *M. marinum*-katushka and treated with MetT as indicated. (B) Quantification of total TUNEL stained cells per granuloma in 5 dpi zebrafish embryos infected with *M. marinum*-katushka and treated with MetT as indicated. (C) Quantification of TUNEL stained cells as a ratio of bacterial content per granuloma in 5 dpi zebrafish embryos treated with MetT as indicated.

### 3. 5. The MetT-induced reduction in bacterial burden is accompanied by an increase in the transcription of inflammatory genes

MetT has been shown to be potently anti-inflammatory, inhibiting the expression of inflammatory cytokines and reducing leukocyte recruitment to sites of active inflammation [21-23]. Given this documented anti-inflammatory effect we investigated the effect that MetT had on the production of inflammatory genes during mycobacterial infection by quantitative real-time PCR (qPCR) at 5 dpi. Transcription of tnfa was unchanged and, surprisingly, there was an increase in the expression of *il1b, il8*, and *mmp9* in MetT-treated infected embryos compared to control embryos (Fig 5A).

**Figure 5:**
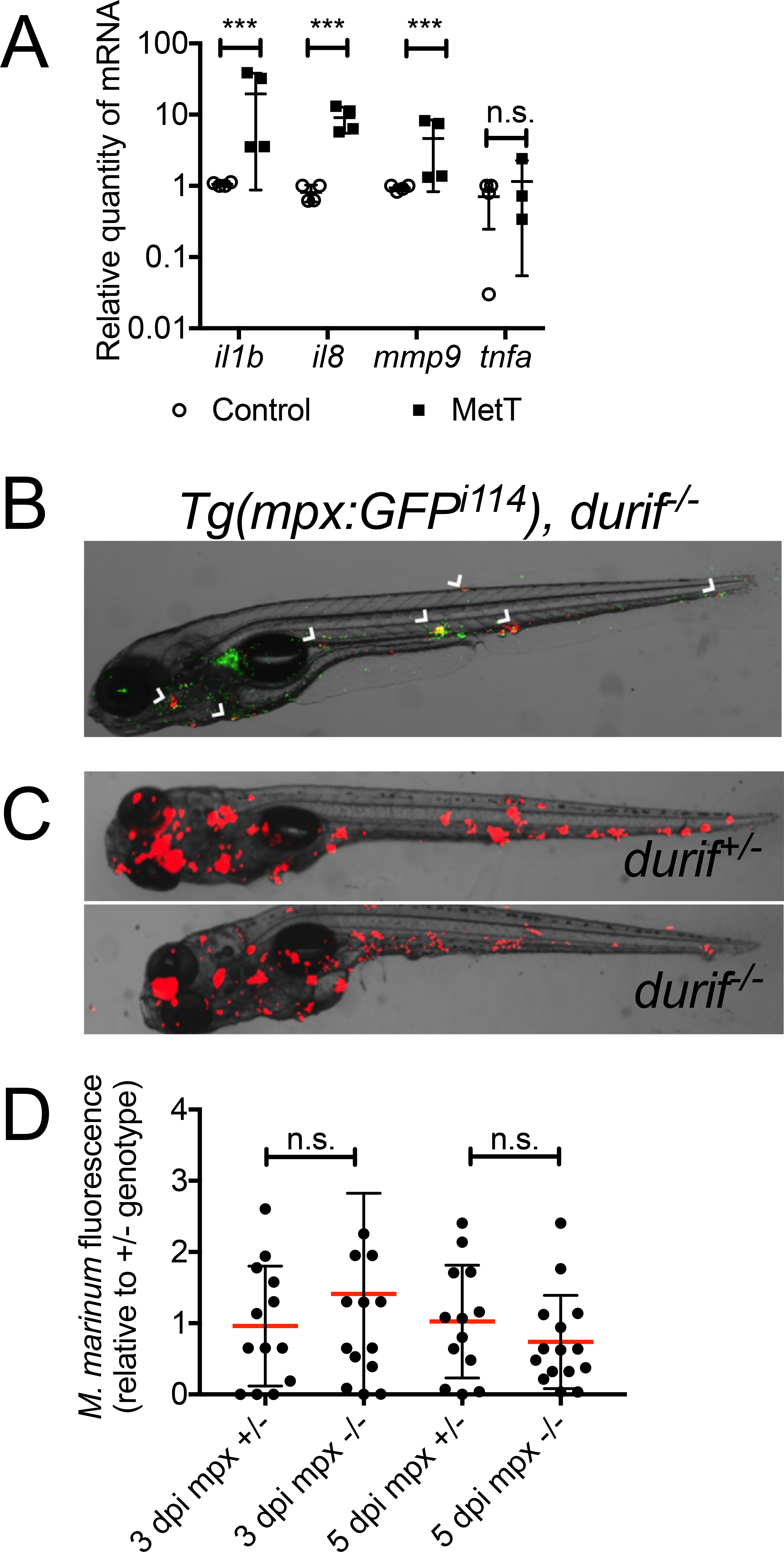
Effects of MetT treatment on innate immune system. (A) qPCR quantification of proinflammatory gene expression in 5 dpi zebrafish embryos infected with *M. marinum* and treated with MetT. (B) Representative image of neutrophil recruitment to granulomas in 5 dpi durif/mpx homozygous mutants infected with *M. marinum*-tomato. White arrowheads indicate granulomas with *mpx* expressing neutrophil recruitment. (C) Representative images of 5 dpi zebrafish *durif* mutant embryos infected with *M. marinum*-tomato. (D) Quantification of bacterial burden by fluorescent pixel count in 5 dpi zebrafish *durif* mutant embryos.

### 3. 6. Neutrophil myeloperoxidase is dispensable for immunity to *M. marinum* infection in zebrafish

Our previous work has demonstrated a potent inhibitor effect of MetT on neutrophil myeloperoxidase [23]. To examine the role of myeloperoxidase in mycobacterial infection, zebrafish embryos were raised from a cross of myeloperoxidase heterozygous knockout adults (mpx +/-) and a myeloperoxidase homozygous knockout (mpx -/-) on a transgenic neutrophil reporting *Tg*(*mpx:EGFP*^*i*114^) background [24]. These embryos were infected with fluorescent *M. marinum*-TdTomato and imaged for bacterial burden at 3 and 5 dpi.

GFP-expressing *mpx* positive neutrophils were present at sites of *M. marinum* infection both in the early granuloma at 3 dpi and at the late granuloma at 5 dpi regardless of functional Mpx state (Fig 5B). There was no gross difference in the pattern of infection, with similar dissemination of the bacteria and size of individual granulomas (Fig 5C). Furthermore, myeloperoxidase knockout animals had similar bacterial burden to their heterozygous clutchmates at 3 and 5 dpi (Fig 5D).

### 3.7. MetT does not inhibit *M. marinum* growth *in vitro* under standard growth conditions

Recent literature has described a direct antimicrobial effect of the NOX and NOS inhibitor diphenyleneiodonium (DPI). Next we determined whether MetT had a similar direct inhibitory effect on *M. marinum* growth that would account for the zebrafish infection phenotypes. Thus, axenic cultures of freshly diluted *M. marinum* cultures were supplemented with MetT at final concentrations of 1 and 5 mM, and colony forming units (CFUs) per mL of culture media were measured. Overall, no significant differences in bacterial CFUs were observed (Fig 6A).

**Figure 6:**
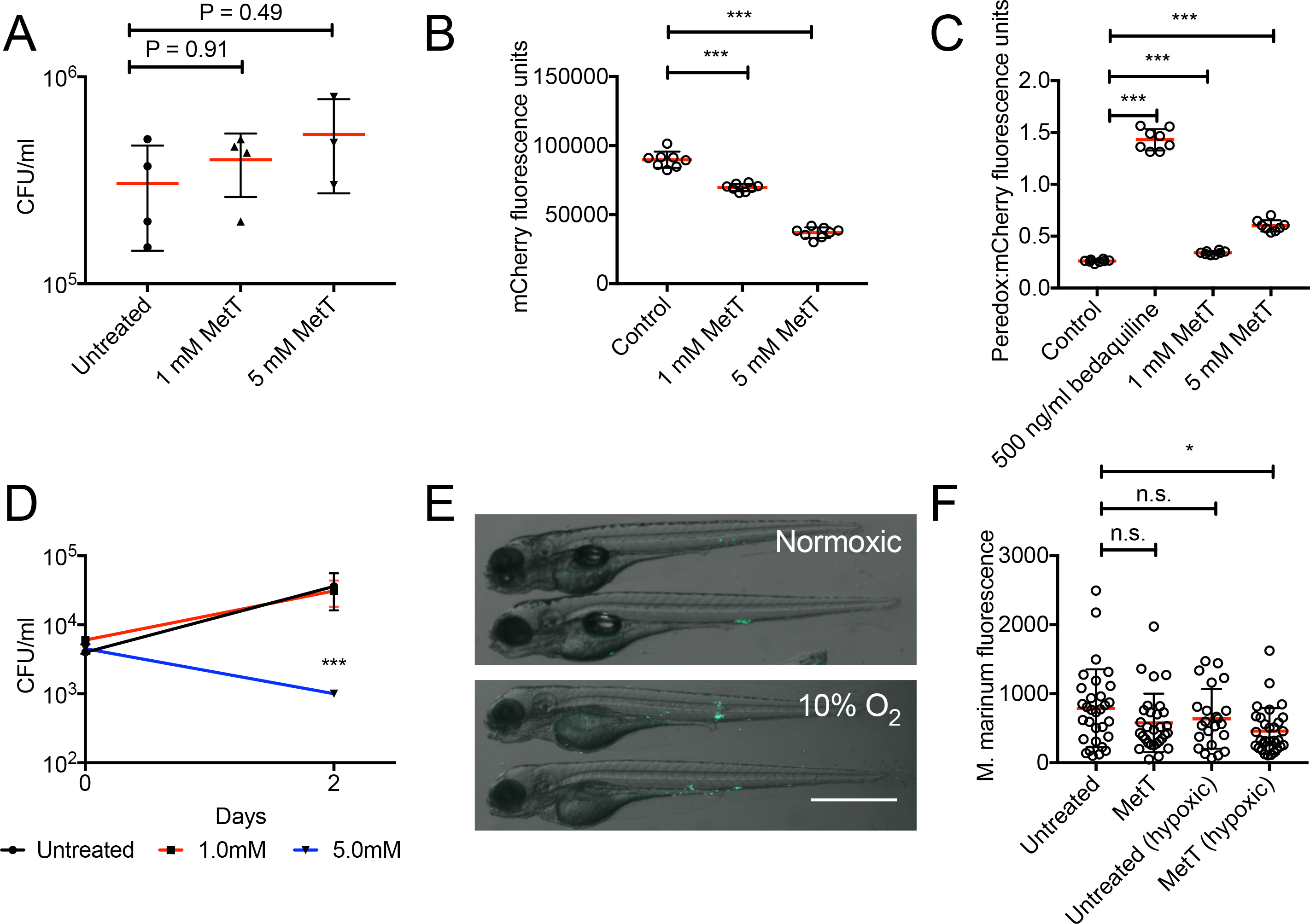
MetT has antibacterial effects against stressed bacteria. (A) CFU recovery from axenic early log phase cultures of *M. marinum* treated with MetT for 2 days. (B) Quantification of constitutive mCherry fluorescence in late log phase cultures of *M. marinum* Δ*esx1* strain carrying pMV762-Peredox-mCherry treated with MetT. (C) Quantification of peredox fluorescence in late log phase cultures of *M. marinum* Δ*esx1* strain carrying pMV762-Peredox-mCherry treated with MetT as an indicator of cellular NADH:NAD build up. (D) CFU recovery from early log phase cultures of *M. marinum* treated with MetT for 2 days (E) Representative images of 5 dpi zebrafish embryos infected with *M. marinum*-wasabi and exposed to 10% O_2_ in a hypoxia chamber. (F) Quantification of bacterial burden by fluorescent pixel count in 4 dpi zebrafish embryos treated MetT and maintained in 10% O_2_ as indicated.

### 3. 8. MetT can disrupt NADH:NAD+ balance in *M. marinum*

Given our observation that MetT affects eukaryotic mitochondrial ROS production, we reasoned MetT may also affect mycobacterial energetics. To visualise disruption of cellular respiration in *M. marinum*, we created a transgenic *M. marinum* Δesx1 strain carrying pMV762-Peredox-mCherry (Addgene 90217) to visualise build up of NADH:NAD ratio [25, 26]. Analysis of constitutive mCherry expression in a late log culture revealed a small but statistically significant decrease in bacteria number following 1 mM MetT treatment and a larger decrease following 5 mM MetT treatment that was correlated with CFU counts (Fig 6B).

Calculation of the ratio of peredox:mCherry fluorescence revealed that 5 mM MetT treatment significantly increased the NADH:NAD ratio, suggesting a mechanism of antibacterial action via electron transport chain disruption, similar to that seen for bedaquiline treatment, in late growth phases *in vitro* (Figure 6C).

### 3. 9. Hypoxic stress sensitises *M. marinum* to MetT *in vitro* and *in vivo*

Hypoxic mycobacteria are more sensitive to electron transport chain disruption and we have previously demonstrated the presence of hypoxia in the zebrafish embryo-*M. marinum* infection model [27-29]. To better model the growth-restrictive conditions found in the *in vivo* infection microenvironment, we next asked if hypoxia could sensitise *M. marinum* to inhibition by MetT *in vitro*. Comparison of recovered CFUs revealed that culture in 5% atmospheric oxygen sensitised *M. marinum* to killing by 5 mM MetT (Fig 6D).

This finding suggested we would be able to increase the *in vivo* sensitivity of *M. marinum* by incubating infected embryos in reduced oxygen conditions. We incubated infected embryos in approximately 10% atmospheric oxygen, which did not affect embryo morphology (Fig 6E). We treated embryos with a shortened suboptimal course of MetT to only 4 dpi, prior to any significant effect on burden, to facilitate identification of an interaction between hypoxia and MetT. We observed an additive effect of hypoxia and MetT treatment *in vivo* (Fig 6F).

## 4. Discussion

We have demonstrated MetT-mediated inhibition of *M. marinum* infection *in vivo* in the zebrafish embryo model, and identified beneficial host- and bacterial-directed effects of MetT treatment. We present a range of live and *in situ* imaging data linking MetT treatment to reduced pathological mitochondrial ROS production and cell death during late mycobacterial infection. Furthermore, stressors found *in vivo* sensitised *M. marinum* to growth restriction or killing by MetT in axenic culture demonstrating a complementary mechanism of host protective action.

The development of multi-drug resistant and extensively drug resistant *Mtb* has shown that the current treatment regimen for tuberculosis is susceptible to evasion and resistance. Traditional pathogen-directed therapies, such as antibiotics, are susceptible to drug resistance caused by spontaneous mutations that are driven by the selective pressure of antibiotic exposure [30]. As host-directed therapies do not directly target bacterial proteins, theoretically there is a reduced risk of drug resistance [31]. However, while there is the reduced risk of drug resistance there is concurrently an increased risk of side-effects since host biology is targeted [32]. While a systematic examination of the toxicity of MetT was beyond the scope of the current study, MetT did not appear to induce an observable pathological phenotype at efficacious doses.

Neutrophils were also examined as a potential target for MetT as they are present at the site of mycobacterial infection and are extensive producers of ROS. Neutrophils primarily kill bacteria through their expression of myeloperoxidase, an enzyme that produces potently microbicidal hypochlorous acid and could trigger macrophage necrosis during infection [8, 33]. Recently, MetT was identified as a potent inhibitor of myeloperoxidase-induced production of hypochlorous acid in an *in vitro* neutrophil co-culture model of acute myocardial infarction [23]. However, using a loss of function mutant of neutrophil myeloperoxidase we show that myeloperoxidase and the subsequent hypochlorous acid is dispensable for control of mycobacterial infection. This finding complements a study by Yang et al. demonstrating that neutrophils have a host protective role in early mycobacterial infection through a NOX dependant pathway and the extensive literature that describes neutrophils having an undefined, but probably limited role in the control of *Mtb* infection in later stages of disease progression [34].

The vast majority of mitochondrial ROS is superoxide generated from the mitochondrial electron transport chain [35]. Cyclic nitroxide derivatives such as MetT have documented superoxide dismutase mimetic effects whereby they reduce superoxide radicals to oxygen and water. It is by this mechanism that we hypothesise MetT reduces mitochondrial oxidants during mycobacterial infection [14, 36, 37]. However, FRR2 is a non-specific oxidant probe, which is fluorescent when oxidised by many reactive oxygen species and nitrogen species. Future studies could confirm whether MetT does indeed reduce superoxide *in vivo*.

Cell death modality is crucial to the pathogenesis of mycobacterial infection [38]. Caspase dependent apoptosis has been demonstrated to be important in the control of mycobacterial infection [39-41]. Pathogenic mycobacteria deploy virulence factors such as ESX1 to supress macrophage apoptosis, suggesting that apoptosis is beneficial to bacterial containment [42]. Conversely necroptosis, a more inflammatory cell death, fuels disease [7, 43]. Given that MetT induces a concurrent reduction in mitochondrial oxidants, cell death and bacterial burden, and that apoptosis is likely host protective, it is an attractive hypothesis that MetT is selectively inhibiting mitochondrial ROS dependant necroptosis. While we demonstrate an inhibition of cell death through TUNEL (see Figure 3.6), TUNEL cannot differentiate between apoptosis and necroptosis [44]. More detailed investigations of cell death are required to determine whether MetT is indeed acting selectively on necroptotic pathways, or rather than as a bulk inhibitor of cell death resulting in the same net effect.

MetT does not inhibit the growth of mycobacteria during the early stages of infection or *in vitro* culture. Instead, inhibition of bacterial growth only becomes apparent around 5 dpi. By 3 dpi, infected macrophages have migrated into deep tissues and have formed tight granuloma aggregations [45]. Between 3 and 5 dpi, there is a dramatic expansion of bacterial load and macrophage cell death. Macrophage depletion around this stage exacerbates necroptosis, and promotes bacterial growth [46]. Our findings support this narrative by MetT-mediated interception of pro-necroptotic ROS production to aid immune containment of infection. From the bacterial perspective, the later timepoint correlates to a much more hostile environment correlating with increased sensitivity to MetT-mediated electron transport chain disruption.

In conclusion, the cyclic nitroxide antioxidant MetT acts on the host and bacterium to inhibit the growth and dissemination of mycobacterial infection in a zebrafish model of tuberculosis. MetT inhibits mitochondrial ROS production and cell death at the site of infection sparing tissue. More broadly, the data presented here demonstrate the value of performing drug discovery in complex model systems to detect *in vivo* microenvironment-drug target interactions.

## Acknowledgements

Dr Kristina Jahn and Sydney Cytometry for assistance with imaging equipment; Garvan Biological Testing Facility staff, Ms Jennifer Brand, Mr Michael Pickering, Ms Rola Bazzi, Dr Lucie Nedved and Dr Stephanie Allison, at the Garvan Institute of Medical Research for maintenance of zebrafish breeding stock; Professor Graham Lieschke for supplying the *durif* mutant line. The authors thank Mr Joshua “Yeezy” Kasparian and Dr Matt Johansen for helpful discussions and technical assistance.

This work was supported by the Australian National Health and Medical Research Council grants APP1099912 and APP1053407; The University of Sydney Fellowship G197581; NSW Ministry of Health under the NSW Health Early-Mid Career Fellowships Scheme H18/31086; and the Kenyon Family Inflammation Award (S.H.O.).

Author contributions: H.B., P.K.W., B.C., and S.H.O. designed the experiments. H.B., W.X., E.H., and S.H.O performed the experiments. S.R. generated the *M. marinum* peredox reporter strain. E.J.N. generated experimental fluorescent redox probes. H.B. and S.O. wrote the paper. W.J.B., P.K.W., B.C., and S.H.O. supervised the project.

